# Multivalent interaction of ESCO2 with the replication machinery is required for cohesion

**DOI:** 10.1101/666115

**Authors:** Dawn Bender, Eulália Maria Lima Da Silva, Jingrong Chen, Annelise Poss, Lauren Gawey, Zane Rulon, Susannah Rankin

**Affiliations:** Program in Cell Cycle and Cancer Biology, Oklahoma Medical Research Foundation, 825 NE 13^th^ St. Oklahoma City, OK 73104; Department of Cell Biology, Oklahoma University Health Science Center, 1100 N Lindsay Ave. Oklahoma City, OK 73104

## Abstract

The tethering together of sister chromatids by the cohesin complex ensures their accurate alignment and segregation during cell division. In vertebrates, the establishment of cohesion between sister chromatids requires the activity of the ESCO2 acetyltransferase, which modifies the Smc3 subunit of cohesin. It was shown recently that ESCO2 promotes cohesion through interaction with the MCM replicative helicase. However, ESCO2 does not significantly colocalize with the MCM helicase, suggesting there may be additional interactions that are important for ESCO2 function. Here we show that ESCO2 is recruited to replication factories, the sites of DNA replication. We show that ESCO2 contains multiple conserved PCNA-interaction motifs in its N-terminus, and that each of these motifs are essential to ESCO2’s ability to establish sister chromatid cohesion. We propose that multiple PCNA interaction motifs embedded in a largely flexible and disordered region of the protein underlie the ability of ESCO2 to establish cohesion between sister chromatids precisely as they are born during DNA replication.

**Summary:** Cohesin modification by the ESCO2 acetyltransferase is required for cohesion between sister chromatids. Here we identify multiple motifs in ESCO2 that mediate its interaction with the replication processivity factor PCNA, and show that their mutation abrogates the ability of ESCO2 to ensure cohesion.

## Introduction

Sister chromatids, the identical products of chromosome replication, are tethered together from the time they are made until cell division when they are segregated into daughter cells. Sister chromatid cohesion is mediated by cohesin, a protein complex that has the capacity to topologically entrap DNA. In addition to its role in tethering sister chromatids together, cohesin also plays critical roles in folding chromosomes into loops and domains throughout interphase, which in turn ensures normal transcription and thus proper development. How cohesin is regulated to result in these very distinct outcomes is not well understood. The activity of the cohesin complex is controlled in part by modulation of the stability of its interaction with chromatin. In higher eukaryotes, cohesin is associated with chromatin throughout interphase, but a small pool becomes more stably bound in a DNA replication-dependent manner, and persists into G2 (Gerlich et al., 2006). This observation and a wealth of genetic and biochemical data suggest that the association of cohesin with chromatin is stabilized by factors or activities associated with DNA replication (Sherwood et al., 2010; Rudra and Skibbens, 2012).

Acetylation of the SMC3 subunit of cohesin by members of the Eco1 family of acetyltransferases stabilizes cohesin binding (Zhang et al., 2008; Sutani et al., 2009; Rolef Ben-Shahar et al., 2008; Unal et al., 2008; Rowland et al., 2009). The founding member of the cohesin acetyltransferase family, the Eco1/Ctf7 protein of *Saccharomyces cerevisiae*, is required for sister chromatid cohesion (Tóth et al., 1999; Skibbens et al., 1999). Vertebrates express two homologs of Eco1, called ESCO1 and ESCO2 (Hou and Zou, 2005; Bellows et al., 2003). In addition to sequences highly conserved with yeast Eco1, both ESCO1 and ESCO2 have N terminal extensions not found in the yeast protein (Fig. 1A). Although ESCO1 and ESCO2 have the same catalytic activity, they make distinct contributions to cohesin regulation. ESCO2 is uniquely able to promote cohesion between sister chromatids, though the majority of SMC3 acetylation is ESCO1 dependent (Song et al., 2012; Alomer et al., 2017).

**Figure 1.**
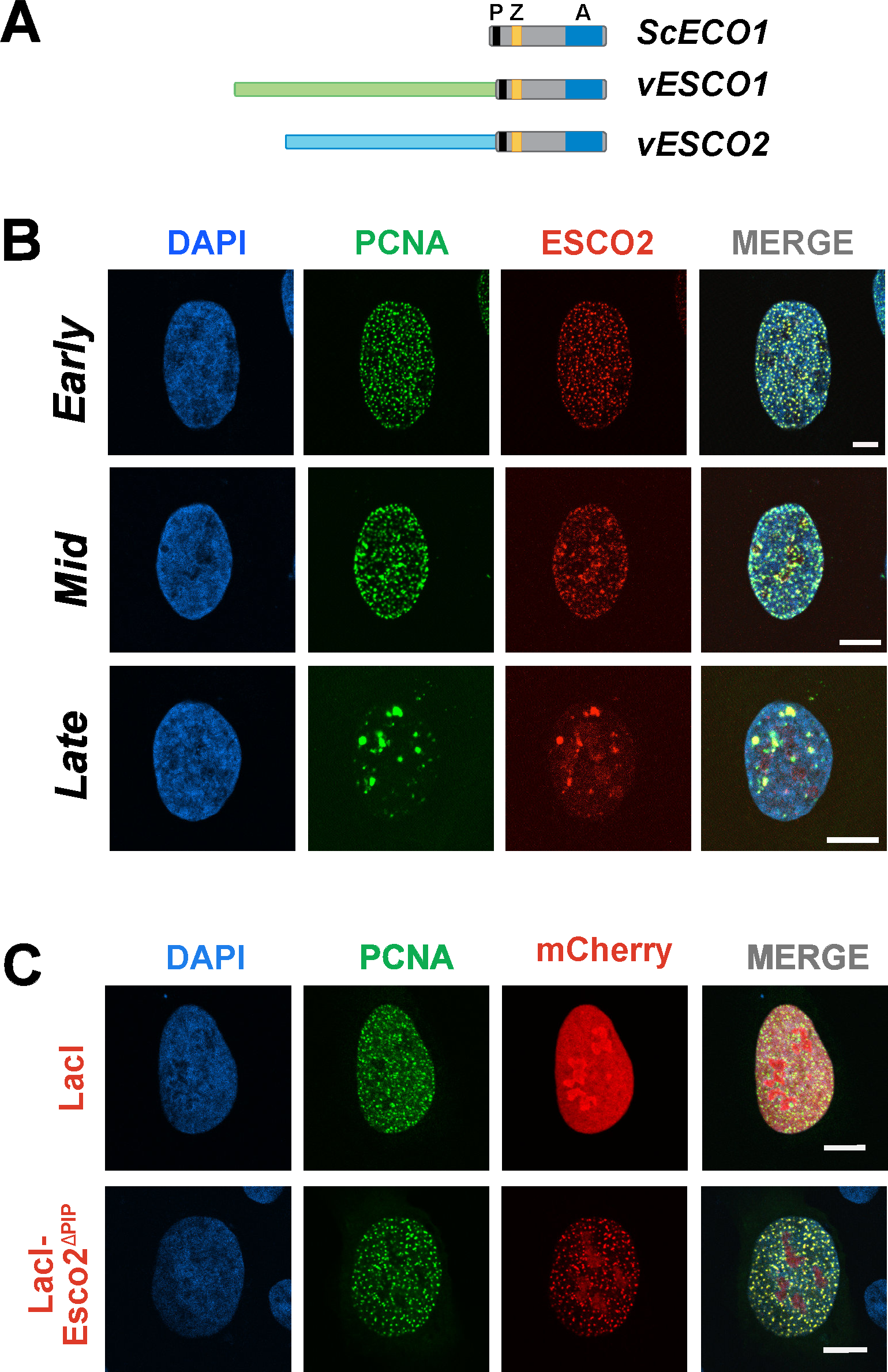
ESCO2 associates with sites of active DNA replication. **A. The Eco acetyltransferases**. Shown are the *Saccharomyces cerevisiae* Eco1 protein and the vertebrate homologs ESCO1 and ESCO2. All three proteins contain a highly conserved domain containing a PIP (<u>P</u>CNA interacting <u>p</u>rotein) box (**P;** black), a zinc finger motif (**Z;** yellow), and the catalytic acetyltransferase domain (**A;** blue). The vertebrate proteins ESCO1 and ESCO2 have unique N-terminal extensions, with no apparent sequence homology to each other (light green and light blue). **B. ESCO2 colocalizes with PCNA in replication foci**. Confocal micrographs of nuclei of U2OS cells co-transfected with mCherry-ESCO2 and GFP-PCNA. ESCO2 colocalized with PCNA in replication foci in patterns of early, mid and late DNA replication. Sites of colocalization appear yellow in the merged images. **C. The PIP box in ESCO2 is dispensable for localization to replication foci**. Confocal micrographs of U2OS cells cotransfected with GFP-PCNA and mCherry-ESCO2 in which the PIP box is deleted. Scale bars: 10μm.

What confers to ESCO2 the unique ability to promote cohesion between sister chromatids? All Eco1 family members contain a conserved PCNA-interacting protein (PIP) box, which in budding yeast has been shown to promote association of Eco1 with the replication factor PCNA, ensuring its association with chromatin during DNA replication (Moldovan et al., 2006). Here we set out to define the elements in ESCO2 that underlie its ability to ensure sister chromatid cohesion. We find that ESCO2 colocalizes with PCNA at sites of active DNA replication, and that it does this independently of the conserved PIP box. We show that ESCO2 interacts with the replication machinery through two non-canonical PCNA-interaction motifs in its N terminal tail. These motifs are embedded in a flexible region of the protein, and the spacing between these motifs varies significantly among ESCO2 proteins from different species. We conclude from these observations that ESCO2 activity is entrained to sites of DNA replication by multiple interactions with the replisome. We propose that the nature of these interactions ensures that ESCO2 retains interaction with this dynamic cellular machine.

## Results

### ESCO2 localizes to sites of DNA replication

We showed previously that ESCO2 function is critically dependent on a conserved PCNA-interacting protein (PIP) box sequence motif near the C terminus of the protein (Fig. 1A), suggesting that the ability of Esco2 to promote cohesion may require direct interaction with PCNA, a critical replication processivity factor (Song et al., 2012). To further explore the potential direct interaction between ESCO2 and PCNA, we co-expressed fluorescent derivatives of both proteins (GFP-PCNA, mCherry-ESCO2) in U2OS cells. Consistent with their direct interaction, the proteins were found colocalized in the nucleus (Fig. 1B). The proteins were found together in “replication foci”, well-documented sites of active DNA replication (Hozák et al., 1994; O’Keefe et al., 1992). During S phase the pattern of replication foci progresses through a stereotypical spatiotemporal program: the pattern begins first with numerous small foci in euchromatic regions in early S phase, followed by accumulation of foci at heterochromatic regions around the nuclear rim in mid S, and finishing with localization to larger patches adjacent to the nucleoli. ESCO2 and PCNA were found colocalized in all of these patterns in fixed cells (Fig. 1B). These data are consistent with interaction of ESCO2 with PCNA at sites of active DNA replication.

The presence of ESCO2 at sites of DNA replication is consistent with the presence of the PIP box, which in fungal models is essential for interaction with chromatin (Moldovan et al., 2006). To test whether ESCO2 colocalizes with the replication machinery through the PIP box motif, we generated a derivative of ESCO2 in which the PIP box was deleted. Unexpectedly, we found that colocalization of PCNA and ESCO2 was unaffected by this mutation: both proteins were found in replication foci, in a manner indistinguishable from the wild type controls (Fig. 1C). We conclude that ESCO2 interacts with the replication machinery independently of the PIP box.

Our data suggested either that sequences other than the PIP box promote ESCO2 interaction with PCNA, or that ESCO2 is recruited to replication foci through proteins other than PCNA. Because co-immunoprecipitation experiments proved inconclusive, we used an alternative approach to test interaction of ESCO2 with PCNA in the nuclear context. To do this, we expressed ESCO2 fused to a fluorescent protein (mCherry) and the lac repressor DNA binding domain in U2OS cells containing an array of *lac* operon operator repeats integrated in the genome. The *lac* repressor DNA-binding domain was recruited to the *lac* operator array, resulting in a bright mCherry focus in the nucleus, as previously (Soutoglou and Misteli, 2008). GFP-labeled PCNA, when coexpressed with the ESCO2 fusion, was recruited to the same foci (Figs. 2A and 2B). Control foci containing mCherry-lacI showed no enrichment for PCNA. A derivative of ESCO2 without the PIP box also recruited PCNA (Fig 2A and 2B). We conclude that ESCO2 interacts with PCNA through motifs other than the canonical PIP box.

**Figure 2.**
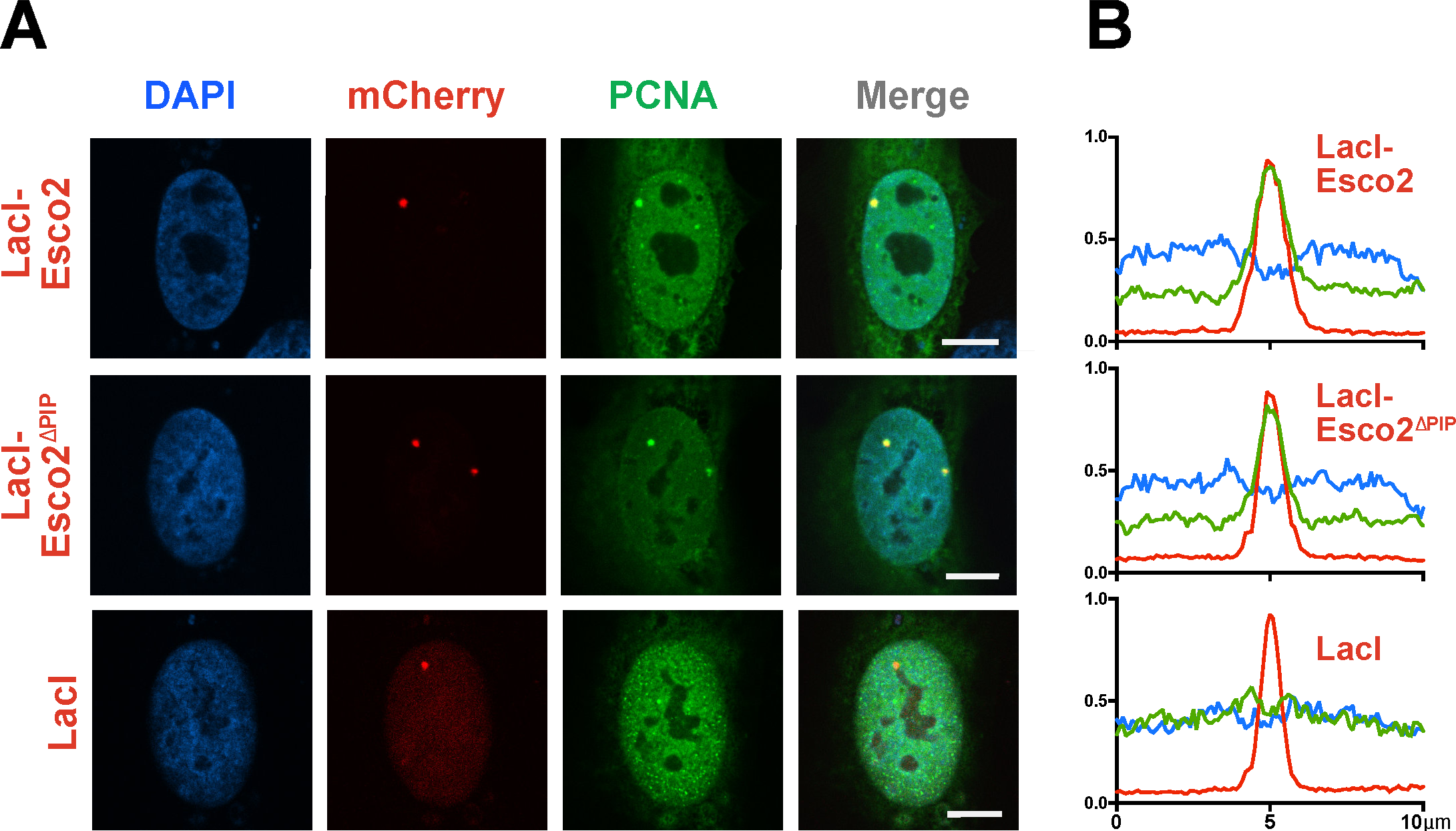
ESCO2 interacts with PCNA outside of replication foci. **A. Tethered ESCO2 recruits PCNA**. An mCherry-lacI-ESCO2 fusion protein, expressed in U2OS cells containing a stably integrated tandem array of lac operator sequences, is recruited to nuclear loci where it can be seen as one or two red foci in confocal micrographs. Cotransfected GFP-PCNA was recruited to the ESCO2 focus (top), and this was unaffected by deletion of the PIP box in ESCO2 (middle). PCNA was not enriched in the foci in cells containing mCherry-lacI (without ESCO2) (bottom). Scale bars: 10μm. **B. Normalized fluorescence intensity**. The fluorescence intensity of lines drawn across the lac operator arrays from a number of cells treated as in **A** was averaged. Fluorescence of mCherry-lacI fusion is indicated by red line, DAPI in blue, and GFP in green. ***n*** >25 cells each sample.

### The ESCO2 N terminus contains multiple essential motifs

Reasoning that sequences important for interaction with the replication machinery are likely to be relatively well conserved, we compared the amino acid sequences of ESCO2 from a number of organisms (Fig. 3A). In general, the N termini are very poorly conserved, with an overall pairwise identity of 26.4%, and only 2.4% identity with the consensus. In contrast, the C termini, starting just after the PIP box, show 72.5% pairwise identity, and 36.1% identity to the consensus. We identified several short sequence elements that were nearly invariant among all of the species analyzed. These included motifs previously shown to be important for chromatin binding (Boxes A and B) (Higashi et al., 2012; Song et al., 2012), as well as a motif near the PIP box, which we call Box C (Fig. 3B).

**Figure 3.**
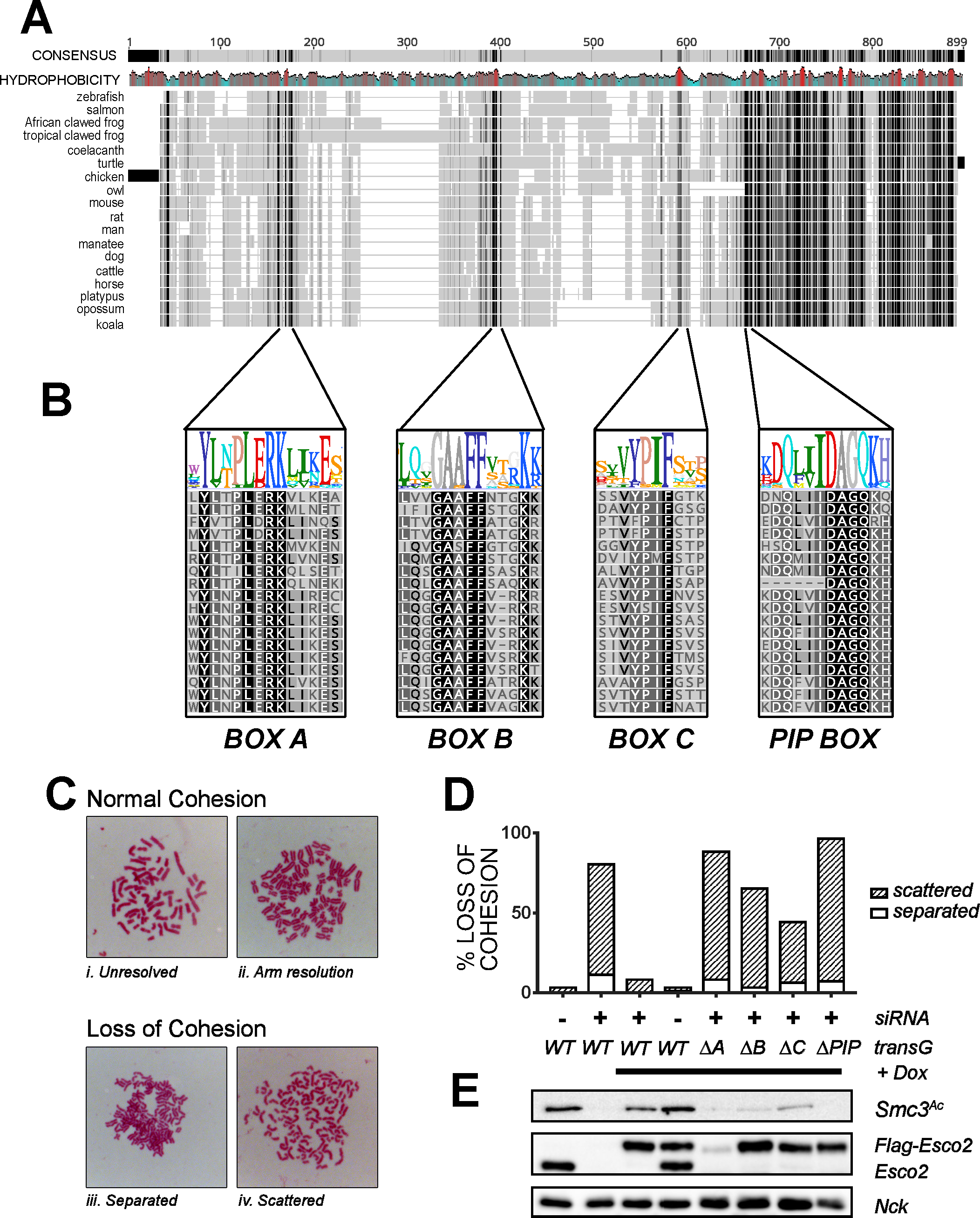
Sequences in the N terminus of ESCO2 are required for cohesion establishment. **A. The N terminus of ESCO2 contains several short, conserved motifs**. Clustal Omega alignment of ESCO2 proteins from the indicated species. The overall consensus is shown at top, in which the grayscale indicates the degree of conservation (black =100% and the lightest gray is<60%), and gaps are shown with a grey line. Mean hydrophobicity is also shown, with red indicating hydrophobic patches. Accession numbers are in Table S1. **B. Enlargement of conserved motifs in the ESCO2 N terminus**. The alignment of the three motifs, Box A, Box B, and Box C, as well as the conserved PIP box, are shown. Sequence logos are colored according to the RasMol scheme (Sayle and Milner-White, 1995). **C. Representative chromosome spreads as analyzed to score sister chromatid cohesion**. Categories **i** and **ii**, in which sister chromatids were clearly tethered together, were considered normal cohesion, while chromatids that were well separated, as in categories **iii** (separated) and **iv** (scattered), were scored as loss of cohesion. **D. Cohesion assay**. HeLa cells expressing siRNA-resistant FLAG-tagged derivatives of ESCO2 with the indicated mutations were treated with siRNA against ESCO2 to deplete endogenous transcripts, and were scored for cohesion as shown in panel C (n≥100/sample). **E. Immunoblot showing expression of ESCO2 transgenes and SMC3 acetylation**. Cell lysates from samples in panel D were probed with antibodies for the indicated proteins. SMC3^Ac^, NCK, and ESCO2 came from the same gel. Nck was used as a loading control. **Tg** = transgene; **Dox** = doxycylin (used to activate expression of transgenes).

The high level of conservation of Boxes A, B, and C suggested that they might be critical for ESCO2 function. To test this, we performed cohesion assays, using a gene knock-down and rescue strategy. HeLa cells lacking a functional ESCO1 gene were engineered to express ESCO2 derivatives as inducible siRNA resistant cDNA transgenes. Expression of endogenous ESCO2 was reduced by siRNA transfection, expression of flag-tagged ESCO2 derivatives was induced by addition of doxycycline, and mitotic spreads were analyzed for cohesion phenotypes (Fig. 3C). We tested each of the conserved motifs, Boxes A, B and C, as well as the conserved PIP box for their impact on cohesion. We found that deletion of any of these conserved motifs resulted in significant loss of cohesion (Fig. 3D). Therefore, each of the conserved motifs is required for full function of ESCO2.

We analyzed the level of SMC3 acetylation in cells expressing the mutant ESCO2 derivatives with deletion of the conserved motifs (Fig. 3E). Because the cells lack functional ESCO1, all acetylation of SMC3 in these samples is ESCO2 dependent (Alomer et al., 2017). We found that in general the relative amount of SMC3 acetylation correlated with the amount of cohesion. For example, cells expressing the PIP box mutant of ESCO2 were strongly compromised in cohesion (>90% loss of cohesion), and had essentially undetectable levels of SMC3 acetylation. In contrast, the Box C mutant was less defective in cohesion establishment, and SMC3 acetylation was reduced but detectable. All of the derivatives of ESCO2 were expressed at similar levels, with the exception of the Box A mutant. The low levels of expression of this mutant may be due to its inability to bind MCM2-7 helicase, as interaction with the MCM helicase has recently been shown to protect ESCO2 from Cul4-DDB1-mediated degradation (Minamino et al., 2018). We conclude from this experiment that all conserved motifs in the otherwise poorly conserved N terminus, Boxes A, B, C, and the PIP box, are required for efficient cohesion establishment or maintenance.

### Box C ensures association with replication foci

The functional difference between ESCO2 and the related ESCO1 maps to their distinct N termini, as opposed to their conserved C terminal acetyltransferase domains (Alomer et al., 2017). This suggests that the N terminus of ESCO2 might be sufficient to promote localization to replication foci. To test this, we fused the ESCO2 N terminus directly to GFP (ESCO2N-GFP), eliminating the catalytic C terminus entirely (Fig. 4A), and co-expressed this fusion together with mRuby-PCNA. We found that ESCO2N-GFP, like the full-length protein, colocalized with PCNA at replication foci (Fig 4B). We then tested derivatives of ESCO2N-GFP in which each of the conserved motifs were deleted. We found that ESCO2N-GFP with deletion of Box A or Box B still colocalized with PCNA at replication foci, but deletion of Box C abrogated this localization (Fig. 4B). We also noted that localization of ESCO2N fusions to the nucleoli, weakly apparent in the full length ESCO2 (Fig. 1B), was more pronounced in the ESCO2N fusions. Partitioning of ESCO2 to nucleoli, while not well understood, has been seen previously, and is consistent with models suggesting that nucleolar function is disrupted in patients with Roberts syndrome, a developmental disorder that results from faulty ESCO2 function (Vega et al., 2005; van der Lelij et al., 2009; Xu et al., 2014).

**Figure 4.**
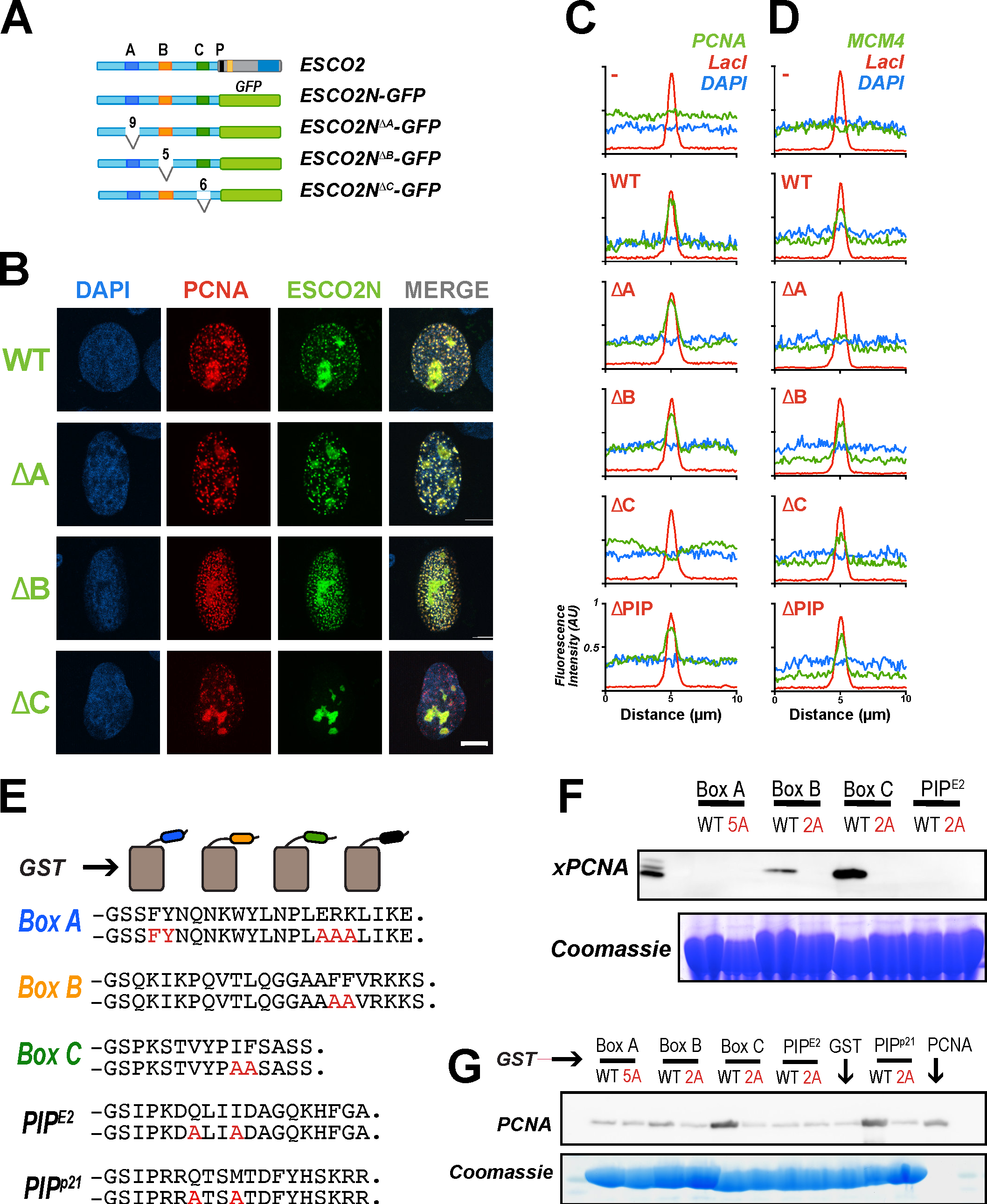
PCNA interacts with ESCO2 motifs *in vivo* and *in vitro*. **A. ESCO2N-GFP fusions**. A cartoon depicting constructs in which the N terminal 375 a.a. of ESCO2 are fused directly to eGFP. Motifs A, B, and C were deleted independently, as shown; numbers indicate the number of amino acids deleted (not drawn to scale). **B. Localization to replication foci**. Confocal images of U2OS cells co-transfected with Ruby-PCNA and the GFP-fusion constructs. Colocalization is indicated in yellow in the merge image, as in Figure 2. **C. Box C is critical for PCNA recruitment by tethered ESCO2**. mCherry-lacI-ESCO2 (full-length) fusions with the deletions indicated in panel **A** were co-expressed with GFP-tagged PCNA as in Figure 2, and the colocalization at nuclear foci was scored by fluorescence intensity profile analysis as in Figure 2. **D. MCM4 recruitment to tethered ESCO2 is dependent upon Box A**. The experiment in Panel C was repeated, only in this case the ESCO2 constructs were co-expressed with mEmerald-MCM4. Recruitment to tethered ESCO2 was scored as in Figure 2. **E. Pull down assay using GST fusion proteins**. Short peptide sequences including Box A, Box B, Box C, or the ESCO2 PIP box motifs were expressed as GST-fusion proteins. A parallel set was made in which alanine substitutions were made at the invariant amino acids (shown in red). The PIP-box from p21 was used as a positive control. **F. Co-precipitation from cell free extracts**. GST fusion proteins shown in panel A were mixed with *Xenopus* egg extract and incubated with glutathione sepharose beads. The beads were washed and bound proteins were eluted and probed for PCNA by immunoblot. A duplicate gel was stained with Coomassie dye to detect the GST fusion proteins. **G. Co-precipitation of purified proteins**. The indicated GST fusion proteins (panel E) were mixed with purified recombinant PCNA, pulled down with glutathione agarose beads and analyzed as in **F** for PCNA.

The experiment in Figure 4B suggested that Box C is essential for interaction with the replication machinery, and is consistent with a model in which ESCO2 interacts, perhaps directly, with PCNA through the Box C motif. To confirm, we again used the assay in which the lacI-ESCO2 is tethered to an integrated array of *lac* operator repeats. Strikingly, the ability of immobilized ESCO2 to recruit PCNA was lost upon deletion of the C box, and unaffected by the absence of Box A, B, or the PIP box (Fig. 4C).

As we were doing this work, two reports were published indicating that Esco2 interacts directly with the MCM2-7 helicase through the Box A motif, and suggesting that this interaction is essential for cohesion establishment (Minamino et al., 2018; Ivanov et al., 2018). We therefore tested our same panel of mutants to determine whether the conserved motifs in ESCO2 are important for interaction with MCM proteins in the tethering assay. Cells with integrated *lac* operator repeats were co-transfected with constructs encoding lacI-ESCO2 and mEmerald-MCM4, and assayed for their colocalization in nuclear foci. While wild-type ESCO2 was able to recruit MCM4, deletion of Box A caused significant loss of interaction in this assay. The other mutants, including deletions of Box B, Box C, or the PIP box were all able to recruit MCM4 (Fig. 4D). We conclude that ESCO2 interacts with both the MCM helicase and with PCNA and that these interactions map to distinct motifs in the ESCO2 N terminus.

### ESCO2 contains multiple PCNA interaction motifs

Although Box A has been shown to promote interaction of ESCO2 with the MCM2-7 helicase, it was still not clear what functional role Boxes B and C play in association with the replication machinery. We noted that the conserved sequences in both Box B and Box C contain invariant pairs of hydrophobic amino acids, FF (Phe-Phe) and IF (Ilu-Phe), respectively. As similar peptides are present in a number of non-canonical PCNA interaction motifs, we tested the possibility that ESCO2 interacts with PCNA through three distinct motifs: Box B, Box C, *and* the PIP box. We expressed short peptides containing the conserved functional motifs as fusions with the glutathione S-transferase protein (GST)(Fig. 4E). The fusion proteins were immobilized on beads, which were then incubated in cell extracts. PCNA bound specifically to the beads carrying GST fusions to both Box B and Box C (Fig. 4B). To our surprise, the previously characterized, functionally critical PIP box at residues 374 to 377 (<u>Q</u>LIIDAGK) did not bind to PCNA in this assay. Similar results were obtained with HeLa nuclear extracts (not shown). PCNA bound most efficiently to the fusion with the Box C peptide, and this binding was abolished when the conserved hydrophobic residues (IF) were changed to alanine (Fig. 4F). Similarly, PCNA bound to the Box B peptide, and this interaction was disrupted by mutation of the FF motif to AA. These data suggest that Box B and Box C are non-canonical PIP boxes.

To understand whether interaction of PCNA with GST fusions was likely to be direct, we purified PCNA and performed co-precipitation experiments with the GST-peptide fusions, including the well-characterized PIP motif from p21 as a positive control. PCNA readily co-precipitated with the Box C peptide, and this interaction was disrupted by mutation of the conserved hydrophobic residues (Fig. 4G). We repeatedly saw weak interaction with the Box B peptide, though this interaction was apparently less robust than that with Box C. The interaction with the Box C peptide was comparable to that of the p21 PIP box. Taken together the experiments in Figures 4F and 4G indicate that ESCO2 contains multiple PCNA interaction motifs in its N terminus. Given that the canonical PIP box is required for full ESCO2 function, we suspect that it may also promote interaction with PCNA, though not in the assays shown here. For simplicity, we will refer to Box B, Box C and the PIP box as PIP1, PIP2, and PIP3 from here forward.

We further analyzed the ESCO2 proteins to better understand how the multiple PIP boxes in ESCO2 might function in context of the full-length protein. We analyzed ESCO2 protein sequences for propensity to adopt specific structures by using the PrDOS algorithm (Ishida and Kinoshita, 2007). This algorithm combines comparison of homologous proteins with amino acid content analysis to predict whether particular sequences are likely to fold into specific structures. Strikingly, all ESCO2 proteins are predicted to be largely disordered throughout their N termini, with clear exceptions at the conserved motifs: Box A, PIP1, PIP2, and PIP3 (Fig. 5A). We found that the spacing between PIP1 and PIP2 varied significantly among species, with a median of 114 amino acids, and a range from 84-164 amino acids, while the spacing between PIP2 and PIP3 was relatively constant (median distance: 47 amino acids, range: 44-56) (Fig. 5B). The distance between Box A and PIP1 was ∼100 amino acids for most species, with a few significant outliers among the species analyzed. We conclude that the essential functional motifs in the ESCO2 N terminus, Box A, PIP1, PIP2, and PIP3, are in a floppy or disordered region of the protein, and that the spacing between these motifs, particularly between PIP1 and PIP2, is not critical to ESCO2 function. This may indicate that the disordered regions simply serve as flexible linkers, consistent with their poor sequence conservation (Fig. 3).

**Figure 5.**
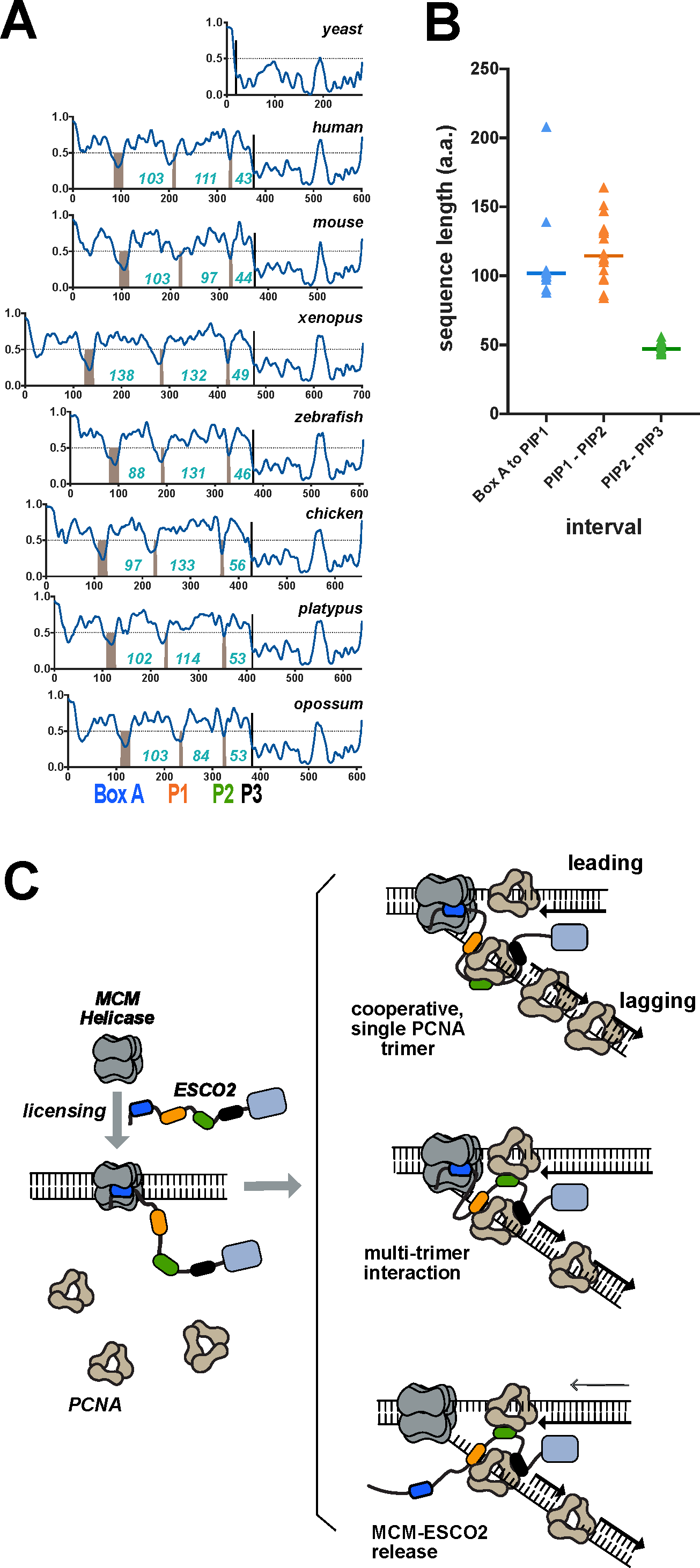
Structural disorder is an intrinsic property of the ESCO2 N terminus. **A. Functional motifs are embedded in disordered region**. The protein sequences of ESCO2 from multiple species were analyzed to detect regions of predicted disorder using the PrDOS algorithm. Sequences above the dotted horizontal line are predicted to be disordered. Conserved motifs, including Box A, PIP1, PIP2 and PIP3 are predicted to be structured, and are indicated by the gray bars. Numbers indicate the number of amino acids between each conserved motif. Accession numbers are shown in Table S2. **B. Spacing between motifs**. The number of amino acids between each of the conserved motifs in the ESCO2 N terminus are indicated in blue. Species included are the same as in A, except that *X. tropicalis* ESCO2 was also included. **C. Models**. ESCO2 is initially recruited to chromatin through interaction with the loaded MCM helicase (gray) through the Box A (blue). Subsequently, ESCO2 associates with sites of active DNA replication through multiple PCNA-interaction motifs (orange, green, and black lozenges). ESCO2 may interact with a single PCNA trimer, as shown at top, or may interact simultaneously with more than one PCNA trimer (middle). The interaction of ESCO2 with MCMs and PCNA may occur simultaneously, or ESCO2 may be released from the MCM complex to associate with PCNA. Multiple, flexible low-affinity interactions may ensure association of ESCO2 with the dynamic replisome

In budding yeast, interaction between PCNA and Eco1 occurs through Eco1’s PIP motif, and this interaction is required for cohesion establishment (Moldovan et al., 2006). Here we show that vertebrate ESCO2, which ensures sister chromatid cohesion, contains two motifs in its N terminus that promote interaction with PCNA. This conclusion is based on colocalization of ESCO2 with PCNA at “replication factories”, the dependence of this interaction on specific sequences in Esco2, and the ability of these same motifs to promote interaction with PCNA in cell free extracts, and with purified proteins. We find that PCNA interacts with each of these peptides in a sequence-specific manner. We have also shown, using a depletion and rescue approach, that each of these motifs is critical to the ability of ESCO2 to promote sister chromatid cohesion. Perhaps surprisingly, although it is essential for cohesion establishment, we were unable to detect a direct interaction between the conserved PIP box in vertebrate ESCO2 (PIP3) and PCNA.

Recent reports show convincingly that ESCO2 is recruited to chromatin by the MCM2-7 helicase and that this interaction is important for cohesin acetylation (Ivanov et al., 2018; Minamino et al., 2018). How then is MCM-dependent recruitment of ESCO2 compatible with the PCNA interaction(s) we report here? The MCM2-7 complex is loaded onto DNA in excess in G1, and only a small fraction of these loaded MCMs are activated to initiate DNA replication (Mahbubani et al., 1997; Walter and Newport, 1997). Indeed, although its activity is required for replication, the MCM helicase is not clearly enriched at sites of active replication nor is it found in replication foci (Aparicio et al., 2012; Krude et al., 1996; Kuipers et al., 2011). It is possible that ESCO2 is initially recruited to chromatin through interaction with the MCM helicase, and subsequently interacts with PCNA during active DNA replication. We do not know currently whether ESCO2 can interact simultaneously with both PCNA and the MCM helicase, or whether these interactions are mutually exclusive. Because PCNA is enriched on chromatin at sites of active DNA replication, we favor a model in which polyvalent interaction of ESCO2 with PCNA ensures SMC3 acetylation specifically at active replication forks, promoting cohesin stabilization precisely when and where sister chromatids are formed.

There is significant degeneracy among functional PIP boxes, and alternative PCNA-interaction motifs have also been identified (De Biasio and Blanco, 2013; Xu et al., 2001; Gilljam et al., 2009). Our data demonstrate that two PCNA interaction motifs, PIP1 and PIP2, mediate interaction between ESCO2 and PCNA. An initial report of the ESCO2-PCNA interaction implicated the PIP3 motif (Moldovan et al., 2006). We tested for, and were unable to detect, this interaction. The previously-reported interaction may have been due to the presence of PIP1 and PIP2 in the region analyzed; a specific requirement for PIP3 was not reported (Moldovan et al., 2006). This is not to say that PIP3 is unimportant; ESCO2 deleted for the PIP3 box localizes normally to replication foci, but is profoundly compromised in cohesion establishment. It is possible that ESCO2 PIP3, which is conserved with the PIP box in *S. cerevisiae* Eco1, in fact promotes interaction with a replication protein other than PCNA. Indeed, a number of PIP-like motifs have been shown to promote such interactions (Boehm and Washington, 2016), and some PIP-like motifs even have dual binding specificity (Iyer et al., 2010). Alternatively, interaction of PIP3 with PCNA may be enhanced in the context of the full-length protein, or seeded by prior PIP2 engagement. PIP1 and PIP2 can be categorized as non-canonical PIP boxes: they contain pairs of hydrophobic amino acids that mediate their interaction with PCNA (Fig. 3).

The interactions of ESCO2 with PCNA may be cooperative, ensuring robust binding through multiple low-affinity interactions. PCNA in its functional state is a ring-shaped homotrimer, in which the flexible interdomain connector loop, where PIP motifs interact, is exposed on the ring surface, and DNA is topologically entrapped within the ring (Dieckman et al., 2012). It is theoretically possible that a single ESCO2 molecule could interact simultaneously with each subunit of a single PCNA trimer, or it may interact with protomers from separate trimers, spanning two or three complexes simultaneously (Fig. 5C). In each of these models, we imagine that interaction with PCNA ensures that ESCO2 is associated with the lagging strand, which is enriched for PCNA, and strongly implicated in cohesion establishment (Shibahara and Stillman, 1999; Rudra and Skibbens, 2012). The presence of multiple interaction motifs is a feature shared with other proteins that associate with PCNA, including Y family DNA polymerases, which participate in translesion synthesis, as well as poly ADP-ribose glycohydrolase (PARG), which is essential for regulation of ADP ribose levels in response to DNA damage (Masuda et al., 2015; Yang, 2014; Kaufmann et al., 2017). ESCO2-PCNA interactions may ensure cohesin acetylation near the single-stranded DNA associated with DNA replication; single-stranded DNA has recently been proposed as an intermediate in cohesion establishment (Murayama et al., 2018).

As in other proteins, the PCNA-interaction motifs in ESCO2 are embedded in an intrinsically disordered region (De Biasio et al., 2014; Kaufmann et al., 2017)(Fig. 5). The disordered nature and variable length of the spacers between the motifs may make an important contribution to how ESCO2 interacts with the replication machinery; the flexibility may reduce the entropic penalty of binding and enhance searching for multiple binding partners. Alternatively, the flexibility may allow ESCO2 to retain association with PCNA as it spins along the DNA helix, through multiple weak and dynamic interactions. Previous attempts to identify ESCO2-interacting proteins using affinity-based methods did not report identification of PCNA, suggesting that the interaction may indeed be weak once removed from chromatin (Minamino et al., 2018; Ivanov et al., 2018). ESCO2 may interact more strongly with PCNA once PCNA is DNA-bound, as reported for other proteins (Havens and Walter, 2009; Gomes and Burgers, 2000).

The presence of multiple binding motifs embedded in a disordered region is consistent with the idea of *allovalency*, in which a single binding site on a receptor (PCNA in this case) can bind several near identical epitopes along the ligand, increasing affinity by increasing the effective local concentration of interaction motifs (Klein et al., 2003). In the context of the PCNA trimer, or indeed the replication fork in which there are multiple PCNA complexes, this kind of interaction may achieve full-on “fuzziness”, in which both partners have multiple interaction sites, and no one particular interaction is favored (Olsen et al., 2017). In “fuzzy” complexes, intrinsically disordered regions retain conformational freedom while interaction motifs are engaged in functional interactions (Tompa and Fuxreiter, 2008). Further experiments will be required to elucidate in detail the nature of the interaction between ESCO2 and the replication machinery.

## Materials and Methods

### Cell culture

HeLa (CCL-2) and U2OS (HTB-96) cells were obtained from ATCC, and U2OS-LacO-I-SceI-TetO cells were obtained from Kerafast (ENH105-FP). All cell lines were cultured in Dulbecco’s Modified Eagles Medium (DMEM, Corning) supplemented with 10% fetal bovine serum (Atlanta Biologicals). Cells were maintained at 37C in a 5% CO_2_ atmosphere. Cells were transfected according to manufacturer’s instructions using Lipofectamine 2000 (Invitrogen) for plasmid DNA or Lipofectamine RNAiMAX (Invitrogen) for siRNA. Stable cell lines with doxycycline-inducible siRNA-resistant Flag tagged ESCO2 cDNAs were generated using HeLa Flp-In T-Rex cells as previously (Alomer et al., 2017). The ESCO2 cDNA was cloned into a pcDNA5/FRT-based Flag-tag vector and co-transfected along with a plasmid expressing the FLP recombinase (pOG44, Invitrogen) using Lipofectamine 2000 (Invitrogen). Cells were selected in 200 μg ml^−1^ Hygromycin B (Gold Biotechnology) and colonies were isolated and expanded, and transgene induction was confirmed by immunoblot.

### Fluorescence analysis

U2OS cells plated on coverslips were transfected with the indicated plasmids, incubated for 24 hours, permeabilized for 5 minutes on ice in pre-permeabilization buffer (20mM HEPES 0.5% Triton X100, 50mM NaCl, 3mM MgCl_2_, 300mM sucrose), then fixed (PBS containing 4% PFA and 0.1% Triton-X100 in) for 15 minutes at RT. Cells were washed with AbDil (20mM TRIS, 150mM NaCl, 2% BSA, 0.1% sodium azide, 0.1% TritonX-100 in PBS) and stained with 4′,6-diamidino-2-phenylindole (DAPI) to label nuclei for one minute at RT and washed again with PBS containing 0.1% TritonX-100. Coverslips were mounted on glass slides using Fluoromount-G (Electron Microscopy Sciences) mounting reagent and sealed with nail polish. Images were collected with a Nikon C2 confocal on a Ti-E motorized inverted microscope using a 60X 1.4 n.a. oil immersion objective lens.

For the tethering assay, U2OS-LacO-I-SceI-TetO cells were plated on coverslips, transfected with indicated plasmids and incubated for 24 hours. Cells on coverslips were fixed (PBS containing 4% PFA and 0.1% Triton-X100) for 15 minutes at RT, washed with AbDil (20mM Tris, 150mM NaCl, 2% BSA, 0.1% sodium azide, 0.1% TritonX-100 in PBS). Cells were stained with DAPI, mounted and imaged as above. NIS Elements software was used to analyze fluorescent intensity along 10μm lines centered on the mCherry focus. Using Prism, the signal for each line scan was normalized from 0 to 1. Graphs depict the mean signal from >25 cells from one representative experiment of three.

### Immunofluorescence

HeLa cells grown on coverslips were fixed with 4% paraformaldehyde in PBS containing 0.1% TritonX-100 at RT for 15 minutes. After fixation, cells were washed and incubated for 30 minutes in AbDil (20mM TRIS pH 7.4, 150mM NaCl, 2% BSA, 0.1% TritonX-100, 0.1% sodium azide) then incubated with appropriate primary antibody diluted in AbDil for 1 hour at RT, washed and probed with secondary for 1 hour at RT. Cells were washed and stained with DAPI to label nucleus for one minute at RT and washed again with TBS with 0.1% TritonX-100. Coverslips were mounted on glass slides and imaged as above.

### Cohesion assays

For cohesion assays, cells with integrated siRNA resistant transgenes were transfected with siRNA against ESCO2 for 4 hours, and cells were left to incubate for 48 hours in doxycycline (2μg/ml). The media was then supplemented with 0.2μg/ml colcemid (AdipoGen) for 15 minutes, and cells were collected with trypsin EDTA, washed with PBS, and treated with hypotonic buffer (0.075M KCl) at 37C for 15 minutes. Cells were collected at 2000G for 5 minutes, resuspended in ∼200 μl of hypotonic buffer, and 5ml of ice-cold fix (3:1 methanol: acetic acid) was added, with gentle mixing. Fixed cells were stored in −20°C. For slide preparation the samples were pelleted and resuspended in freshly prepared fix, then dropped onto slides, steamed over a hot water bath for 15 seconds, and dried on a slide warmer at 60C. The slides were stained with Giemsa (VWR) and coverslips were mounted with Permount (Fisher). Images were collected using a Zeiss Axio Imager Z.1, using 63X 1.4 n.a. oil immersion lens. Phenotypes were assigned to mitotic chromosome spreads as previously (Alomer et al., 2017). Samples were scored blind and each experiment was repeated at least 3 times.

### Co-precipitation assays

GST-peptide fusions were purified in Lysis Buffer (50mM TRIS 8.2, 500mM KCl, 1 mM DTT, 1% Triton X-100) and left on Glutathione Sepharose 4B (GE Healthcare) beads following purification. The beads were washed extensively in nuclear extract buffer (20mM HEPES 7.9, 100mM KCl, 0.2 mM EDTA, 20%glycerol, 2mM DTT). The beads were then incubated with *Xenopus* egg extract (Murray, 1991), and mixed on a twirler at 4C for 1-2 hours. The beads were washed three times in NEB, and bead-associated proteins were eluted by boiling the beads in sample buffer. To analyze the interactions between purified proteins, purified GST fusions where incubated with beads for 1 hour at 4C. Bacterially expressed 6His-GFP-PCNA (>95% purity) was added and allowed to incubate for 2 hours on a twirler at 4C. Beads were washed three times with low salt buffer (50mM Tris, 100mM NaCl, 10mM MgCl_2_, 0.05% Tween-20) prior to elution by boiling in SDS-PAGE sample buffer. Proteins eluted from the beads were analyzed by immunoblot and Coomassie stain.

### Gels and immunoblots

For immunoblots, protein samples were resolved on 7–15% gradient SDS-PAGE gels, transferred to nitrocellulose membranes, incubated with 5% milk in TRIS-buffered saline (TBS), and probed with empirically determined concentrations of primary antibodies overnight at 4°C. Horseradish peroxidase-labeled secondary antibodies were detected with chemiluminescent substrate (Licor Biosciences) and signals were collected using an Azure C600 CCD imager (Azure Biosystems).

### Protein sequence analysis

Protein alignments (Fig. 3) were done using Geneious 2019.0 (Biomatters) using the built-in Clustal Omega algorithm and default parameters. The sequence logos were generated in Geneious using Rasmol colors. The hydrophobicity prediction graph was generated using a sliding 5 a.a. window.

### Plasmids

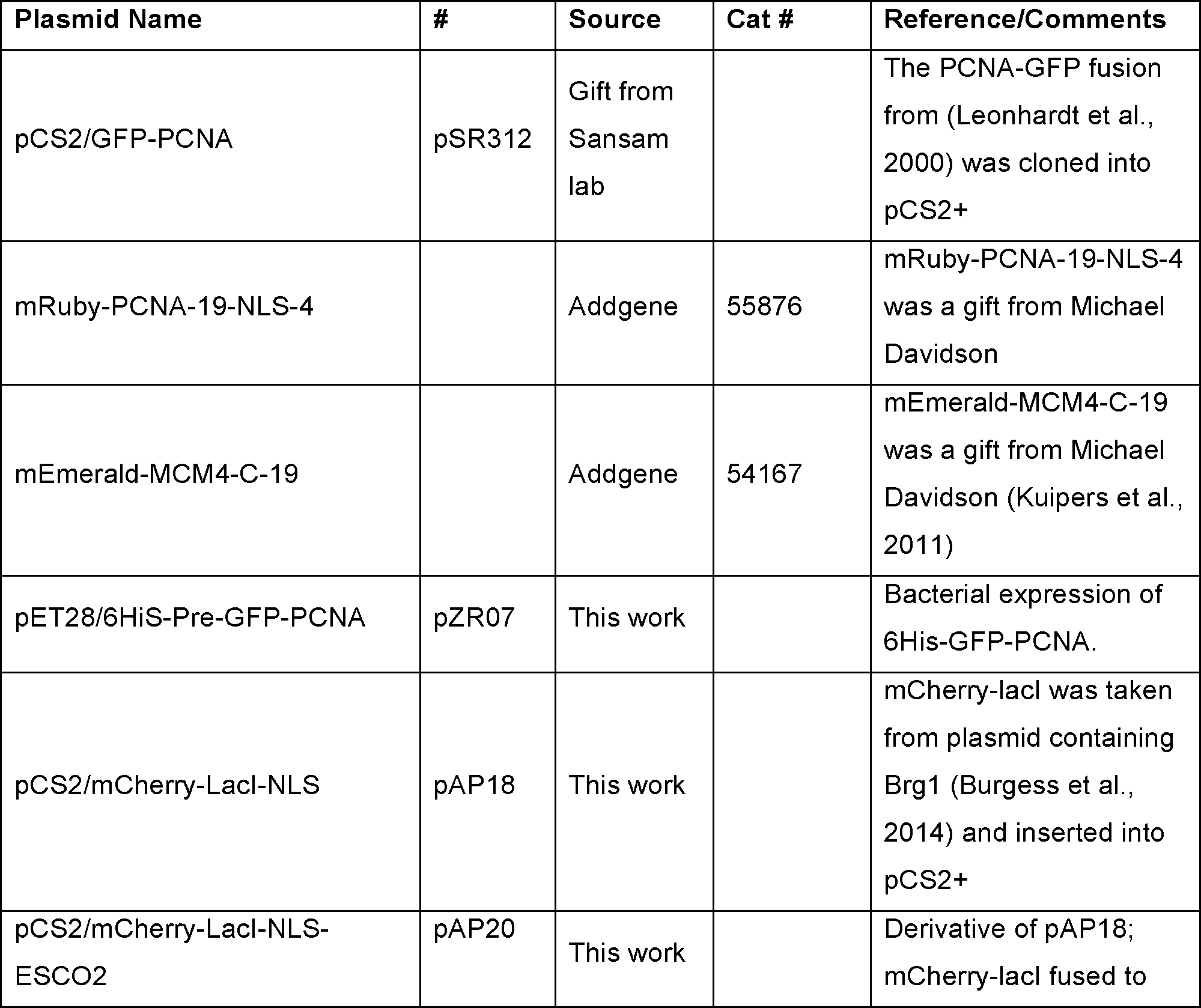
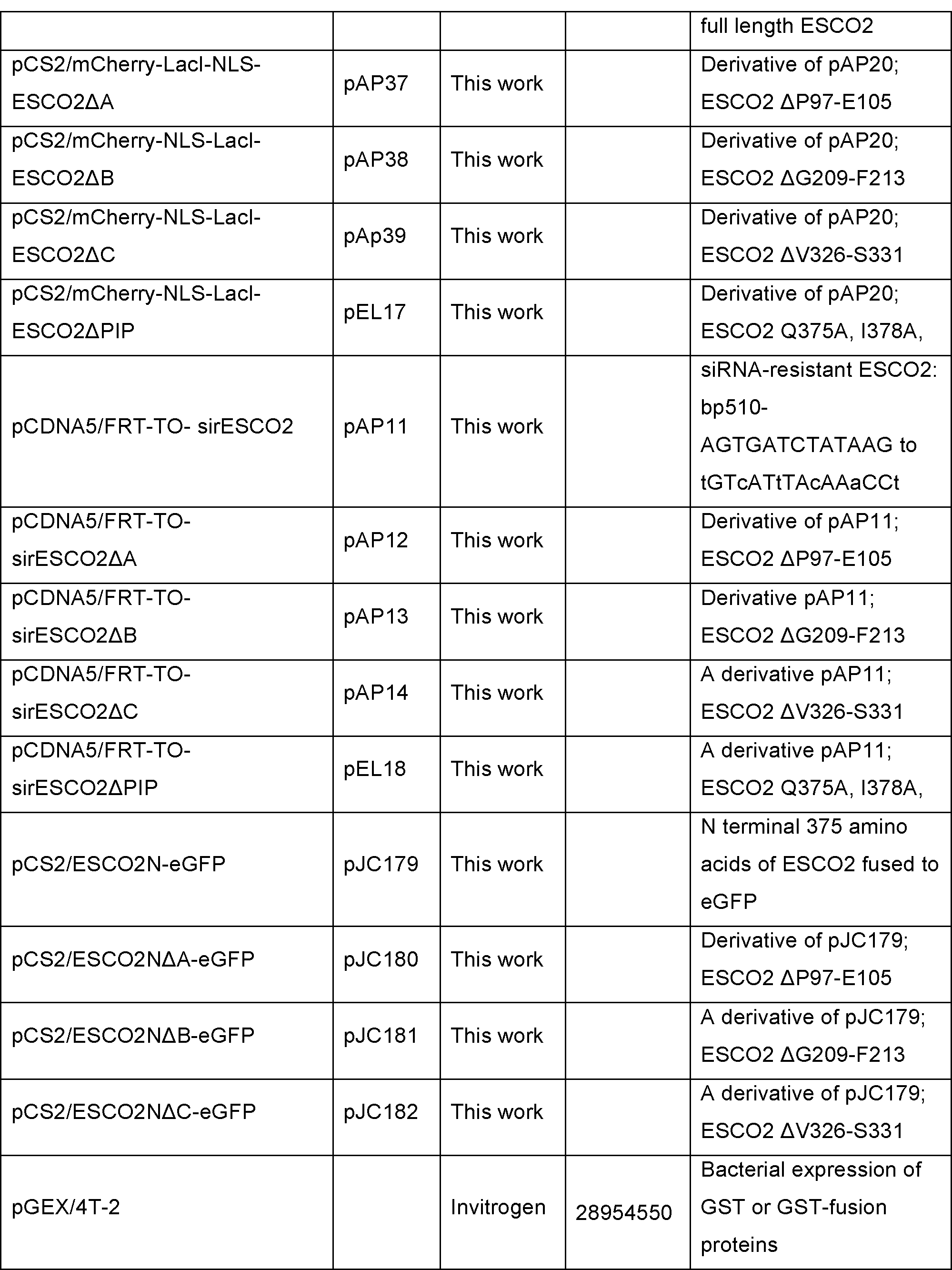
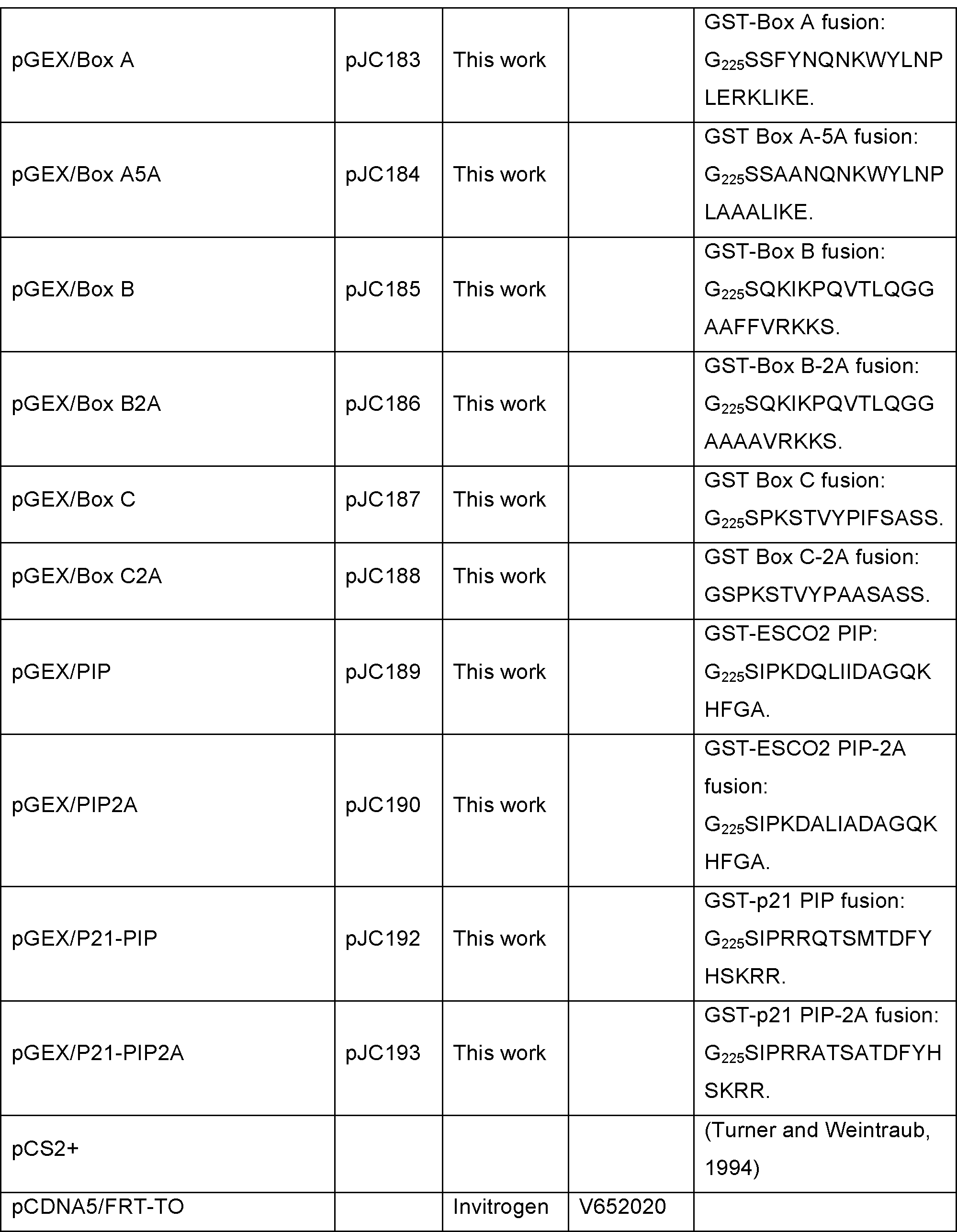

### siRNA

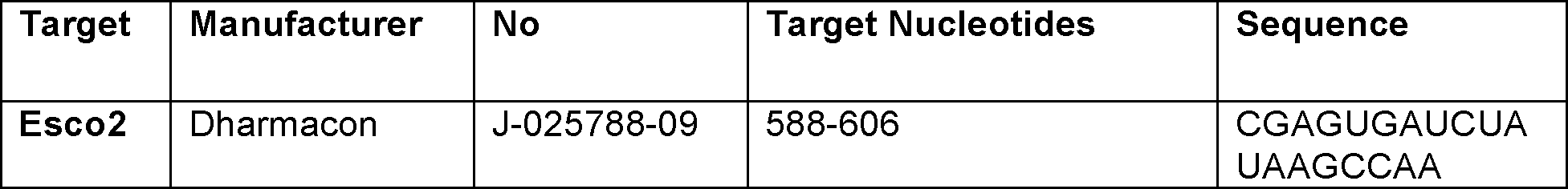

### Antibodies

| Name | Source | No | Lot | Concentration |
| --- | --- | --- | --- | --- |
| PCNA (FL-261) | Santa Cruz | Sc-7907 | F1711 | IF 1:500 |
| MCM4 | Bethyl Labs | A300-193A |  | IF 1:500 |
| SMC3<sup>Ac</sup> | Gift from<br>Katsuhiko Shirahige<br>(Nishiyama et al.,<br>2010) |  |  | IB 1:1500 |
| SMC3 | (Lafont et al., 2010) |  |  |  |
| NCK | NeoMarkers | RB0024 |  | IB 1:1000 |
| ESCO2 | Bethyl Labs | A301-689A | multiple | IB 1:1000 |
| HA-Tag | Cell Signaling | C29F4 | 8 | IF 1:500 |
| Anti-Flag | Sigma | F1804 | 101M6216 | 1:1000 |

## Acknowledgements

This work was supported by NIH grant R01GM101250, and the Oklahoma Center for Adult Stem Cell Research, both to SR, a Pat and Don Capra Predoctoral Fellowship to DEB, and a fellowship from the National Council for Scientific and Technological Development of Brazil to EMLdS. We are grateful to Dean Dawson for careful review of the manuscript and to all members of the Program in Cell Cycle and Cancer Biology for helpful discussion during the course of this work.

---

**Table S1.** Accession numbers from alignment in Figure 3A.

| Accession | organism | common name | Length (a.a.) |
| --- | --- | --- | --- |
| NP_001003872 | Danio rerio | zebrafish | 609 |
| NP_001133441 | Salmo salar | salmon | 631 |
| NP_001089603 | Xenopus laevis | African clawed frog | 702 |
| NP_001121462 | Xenopus tropicalis | tropical clawed frog | 770 |
| XP_014343155 | Latimeria chalumnae | coelacanth | 702 |
| XP_006132649 | Pelodiscus sinensis | turtle | 685 |
| XP_420012 | Gallus gallus | chicken | 691 |
| XP_026702143 | Athene cunicularia | owl | 624 |
| NP_082315 | Mus musculus | mouse | 592 |
| XP_006252234 | Rattus norvegicus | rat | 595 |
| XP_011542723 | Homo sapiens | man | 601 |
| XP_004382239 | Trichechus manatus latirostris | manatee | 614 |
| XP_022265548 | Canis lupus familiaris | dog | 598 |
| NP_001094652 | Bos taurus | cattle | 610 |
| XP_014593313 | Equus caballus | horse | 612 |
| XP_007664914 | Ornithorhynchus anatinus | platypus | 639 |
| XP_007475966 | Monodelphis domestica | opossum | 610 |
| XP_020839651 | Phascolarctos cinereus | koala | 607 |

**Table S2.**
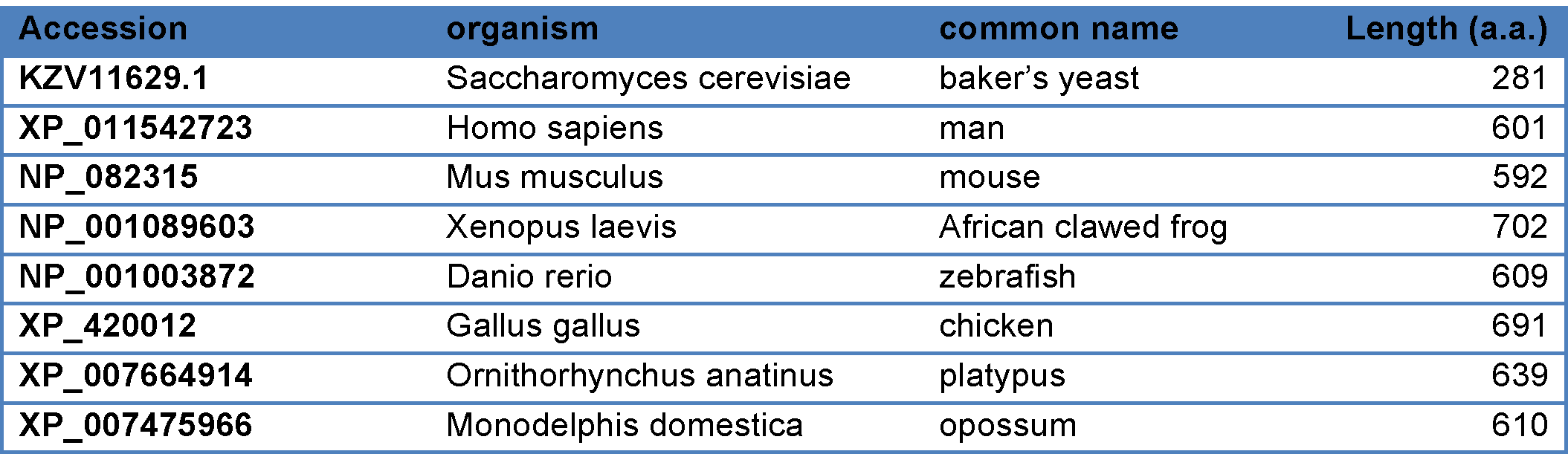
Accession numbers of sequences analyzed in Figure 5A.

